# Application of nanotags and nanobodies for live cell single-molecule imaging of the Z-ring in *Escherichia coli*

**DOI:** 10.1101/2023.02.19.529144

**Authors:** Emma Westlund, Axel Bergenstråle, Alaska Pokhrel, Helena Chan, Ulf Skoglund, Daniel O. Daley, Bill Söderström

**Affiliations:** Australian Institute for Microbiology and Infection, University of Technology Sydney, Ultimo, NSW, 2007 Australia; Department of Biochemistry and Biophysics, Stockholm University, SE-106 91 Stockholm, Sweden; Structural Cellular Biology Unit, Okinawa Institute of Science and Technology, 904-0495 Okinawa, Japan

## Abstract

Understanding where proteins are localized in a bacterial cell is essential for understanding their function and regulation. This is particularly important for proteins that are involved in cell division, which localize at the division septum and assemble into highly regulated complexes. Current knowledge of these complexes has been greatly facilitated by super-resolution imaging using fluorescent protein fusions. Herein we demonstrate with FtsZ that single-molecule PALM images can be obtained *in-vivo* using a genetically fused nanotag (ALFA) and a corresponding nanobody fused to mEos3.2. The methodology presented is applicable to other bacterial proteins.

## Introduction

In recent years super-resolution fluorescence microscopy methods have taken centre stage in microbial cell biology as they can resolve cellular structures well below the diffraction limit (Xiao and Dufrene 2016). One cellular system that has benefitted greatly from these methods is the bacterial cell division machinery or the ‘divisome’. The ‘divisome’ is a complex and transient macromolecular protein assembly, containing >35 different proteins that span all compartments of the cell (Attaibi and den Blaauwen 2022, Haeusser and Margolin 2016). First to arrive at the division site is FtsZ (McQuillen and Xiao 2020), a protein that functions as a molecular foreman that guides the assembly and dynamics of other divisome proteins (den Blaauwen, et al. 2017, Erickson, et al. 2010). Due to its broad conservation over many bacterial species, FtsZ could also be a potent target for new antibiotics (Kusuma, et al. 2019). As such an interest in understanding the function of FtsZ has driven the cell division field for the last 25 years (den Blaauwen, et al. 2017).

One technical aspect that has hampered studies of FtsZ and other cell division proteins is that reliable labelling is an ongoing challenge for fluorescence microscopy imaging, especially in super-resolution imaging. Considerable progress has been made using N- and C-terminal fusions to fluorescent proteins (FPs), which provide a multi-wavelength tool-box for all modes of fluorescence imaging techniques (Holden 2018, Xiao and Dufrene 2016). And since FP’s can be imaged in live cells, temporal changes in localization can be tracked (Holden 2018, Rowlett and Margolin 2014, Strauss, et al. 2012). A major downside to FP fusions is that they are large (~25 kDa) and that they can perturb the function of the protein they are fused to (as judged by the fact that they cannot always fully complement the phenotype of a deletion strain).

As such they often need to be expressed from plasmids and studied in the context of the native (chromosomally encoded) division protein. An unwanted side-effect of doing this is that the amount of protein in the cell is higher than normal, which can perturb the timing of division and result in an elongated cell phenotype. Eriksson and colleagues elegantly worked around this problem, by exploring internal fusion points for FPs in FtsZ (Moore, et al. 2017). They noted that all N- and C-terminal fusions are not functional, but that position G55:Q56 could functionally tolerate some but not all FPs. Recently similar approaches have been applied to other division proteins, e.g. FtsN (Lyu, et al. 2022) and FtsA (Cameron and Margolin 2023). It should be noted that this approach is extremely time-consuming and not broadly applicable.

An alternative approach is to use fluorescently labelled antibodies to label the native protein. Here a primary antibody binds to the protein of interest, and a fluorescent dye labelled secondary antibody is used to label the primary antibody. This approach, immunofluorescence microscopy or IFM, has been widely used but is limited by the availability and quality of antisera. Another major drawback here is the large linkage error, defined as the distance from target protein to fluorescent label (Lelek, et al. 2021). The linkage error can be on the order of 50 - 30 nm as antibodies are relatively large and thus of comparable size to the resolving power of various super-resolution microscopy methodologies. This problem is compounded by the fact that most antibodies are polyclonal and bind to multiple epitopes on a protein of interest. These drawbacks create uncertainty about the localization of the protein being studied, which is counter-productive as super resolution imaging system can have practical resolving powers down to a few tens of nanometers (Carrington, et al. 2019). One way of circumventing this issue is to use alternative protein tags, e.g. the Halo-tag (Los, et al. 2008). These tags are fused to your target protein, whereby a fluorescently labelled substrate (typically an organic dye) is added to bind to the tag. The halo tag has been successfully used in bacterial cell division studies previously and is especially well suited for single-molecule tracking type of experiments (McCausland, et al. 2021). One big drawback, however, is that the Halo-tag itself is relatively large (comparable to the size of a FP).

Herein have we instead explored the use of nanotags (NTs) and fluorescently labelled nanobodies (NBs) for labelling division proteins in *E. coli*. Specifically, we have used the ALFA-tag (Gotzke, et al. 2019), a small and highly versatile protein tag that can be used for a wide range of biological applications (Akhuli, et al. 2022). In bacteriology, the ALFA-tag has thus far been utilised in super-resolution microscopy of DNA damage repair as well as in cell wall synthesis using structure biology approaches (Shlosman, et al. 2022, Wiktor, et al. 2021), but to our knowledge never before in a dynamic live cell setting. We demonstrate that the ALFA-tag can be integrated into the genome of *E. coli*, without affecting the function of the essential cell division protein FtsZ. We further demonstrate that ALFA-labelled FtsZ can be imaged by both STimulated Emission Depletion (STED) using fluorescently labelled NBs in an immunofluorescence approach, and importantly by expressing fluorescently labelled NBs from plasmids using live cell PhotoActivated Localization Microscopy (PALM). The *in vivo* approach has a number of advantages over regular FPs fusions and immunofluorescence microscopy, which are discussed.

## Results

Our goals in this study were (1) to determine whether an essential, and normally difficult to tag *E. coli* cell division protein, would remain functional when fused to a NT, and (2) if it could be imaged using super-resolution microscopy. The ALFA-tag is 39 nucleotides long and encodes a 13 amino acid peptide that forms an a-helix (Figure 1A) (Gotzke, et al. 2019). The coding sequence for ALFA was successfully inserted into the chromosomal copy of *ftsZ* in the *E. coli* strain MG1655 using CRISPR Optimized MAGE Recombineering (CRMAGE) (Ronda, et al. 2016). It was possible to integrate the coding sequence for ALFA in the region of *ftsZ* corresponding to G55:Q56 by using flanking sequences (GSTLE and LEGST) (Figure 1B). Attempts to insert the coding sequence for ALFA at the same position (G55:Q56) but without the flanking sequences, or at the 3’ end of FtsZ were not successful, suggesting that the designs were not functionally tolerated (as noted previously with FPs (Moore, et al. 2017)). The strain, herein denoted FtsZ-ALFA had no visible phenotype when visualized by light microscopy (Figure 1C). The cell lengths were indistinguishable from the wildtype parent strain (Figure 1D), as was the growth curve (Figure 1E). Quantitative Western blotting of cells harvested in the exponential phase indicated that the expression levels of FtsZ in wild type and the FtsZ-ALFA strain were comparable (Figure 1F). Taken together, these data indicate that ALFA can be functionally fused to FtsZ in a non-disruptive way. To determine if this tagging strategy was suitable for super resolution imaging, we used two orthogonal labelling approaches: Immunofluorescence labelling in fixed cells (Figure 1G), and plasmid expression of α-ALFA fused to mEos3.2 (a photoconvertible FP) in live cells (Figure 1H). We then visualised the tagged proteins with two standard super-resolution techniques: STimulated Emission Depletion (STED) (Hein, et al. 2008, Vicidomini, et al. 2011) and single-molecule PhotoActivated Localization Microscopy (PALM) (Betzig, et al. 2006, Greenfield, et al. 2009).

**Figure 1.**
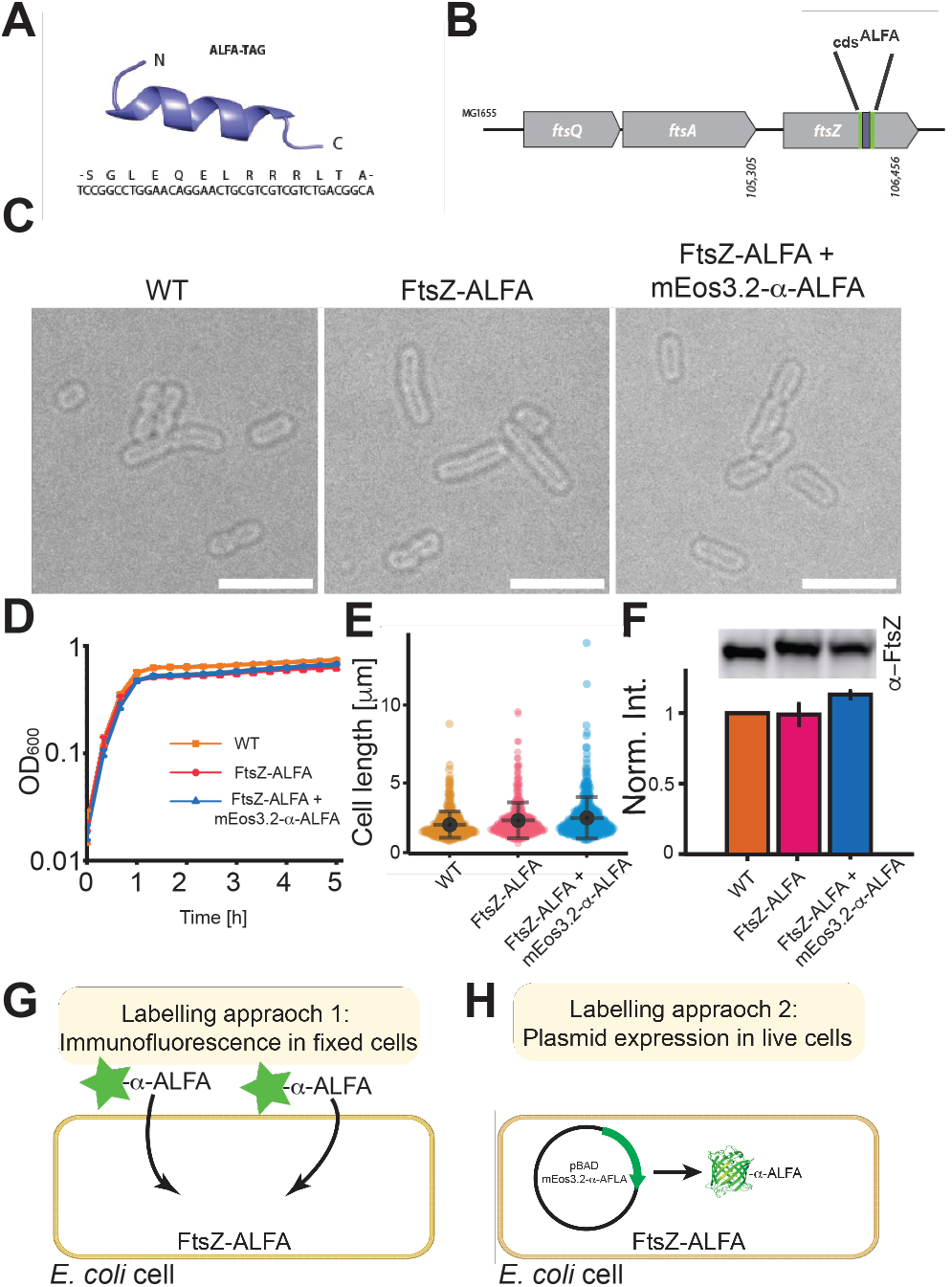
Cell viability of engineered FtsZ-ALFA strain and labelling approaches. **A**, Structure and sequence of the ALFA-tag (Gotzke, et al. 2019). **B**, Locus on the chromosome in strain MG1655 where the ALFA-tag is incorporated. **C**, Bright field images of strain used in this study. No morphological differences are noticeable. Scale bars = 5 μm. **D**, Growth curves of strains used (n = 3 for each strain). **E**, Cell length measurements. WT = 2.34 ± 0.83 μm, FtsZ-ALFA = 2.61 ±1.16 μm, FtsZ-ALFA + mEos3.2-α-ALFA = 2.78 ± 1.33 μm (n = 214 for each strain). Mean ± S.D. **F**, Quantitative Western blotting indicated that neither the ALFA-tag nor the additional expression of mEos3.2-α-ALFA altered the overall expression of FtsZ to any larger degree. WT = 1, FtsZ-ALFA = 0.99 ± 0.08, FtsZ-ALFA + mEos3.2-α-ALFA = 1.13 ± 0.03. Mean ± S.D. **G**, Approach 1: immunofluorescence labelling. Cells were fixed, membranes permeabilized followed by labelling with ATTO488 tagged α-ALFA nanobodies recognizing the ALFA-tag. **H**, Approach 2: Plasmid expression in live cells. Cells were transformed with a plasmid encoding for mEos3.2-α-ALFA nanobody. In both approaches FtsZ-ALFA is labelled with fluorescently tagged nanobodies and imaged using super-resolution approaches (Figure 2 and 4). STED and immunofluorescence

As a first approach, we used an established immunofluorescence methodology to label FtsZ-ALFA. Cells were fixed with formaldehyde, membranes were permeabilized using standard protocols, then FtsZ-ALFA was labelled with α-ALFA-ATTO488 nanobodies and imaged by super-resolution STED (Vicidomini, et al. 2013). As anticipated most elongated cells had a visible FtsZ band at the future division site (Figure 2A) (Sun and Margolin 1998). Images and quantitative measurements of the Full Width at Half Maxima (FWHM) indicated that the resolution improvement by STED over conventional confocal imaging was approximately three-fold (Figure 2B). The average width of the FtsZ-ALFA bands observed from the STED images was determined to be 94 ±12 nm compared to 243 ±26 nm for confocal (Figure 2C). These STED values were slightly lower than previously reported dimensions found using STED on IFM labelled FtsZ in *E. coli* (~110 nm) (Söderström, et al. 2018b), possibly since the linkage error was reduced using the nanotag approach. FtsZ forms filaments that treadmill around the division septum (Yang, et al. 2017). Under the super-resolution light microscope these FtsZ filaments resemble elongated clusters (Fu, et al. 2010, Lyu, et al. 2022, Söderström, et al. 2018b). Here we visualised FtsZ-ALFA rings in cells trapped in a vertical position using a micron-cage approach (Figure 2D) (Söderström, et al. 2018b). As expected, clearly resolvable FtsZ-ALFA clusters were distributed around the division and a pseudo-timecourse could be captured in constricting cells with radii below 100 nm (Figure 2E).

**Figure 2.**
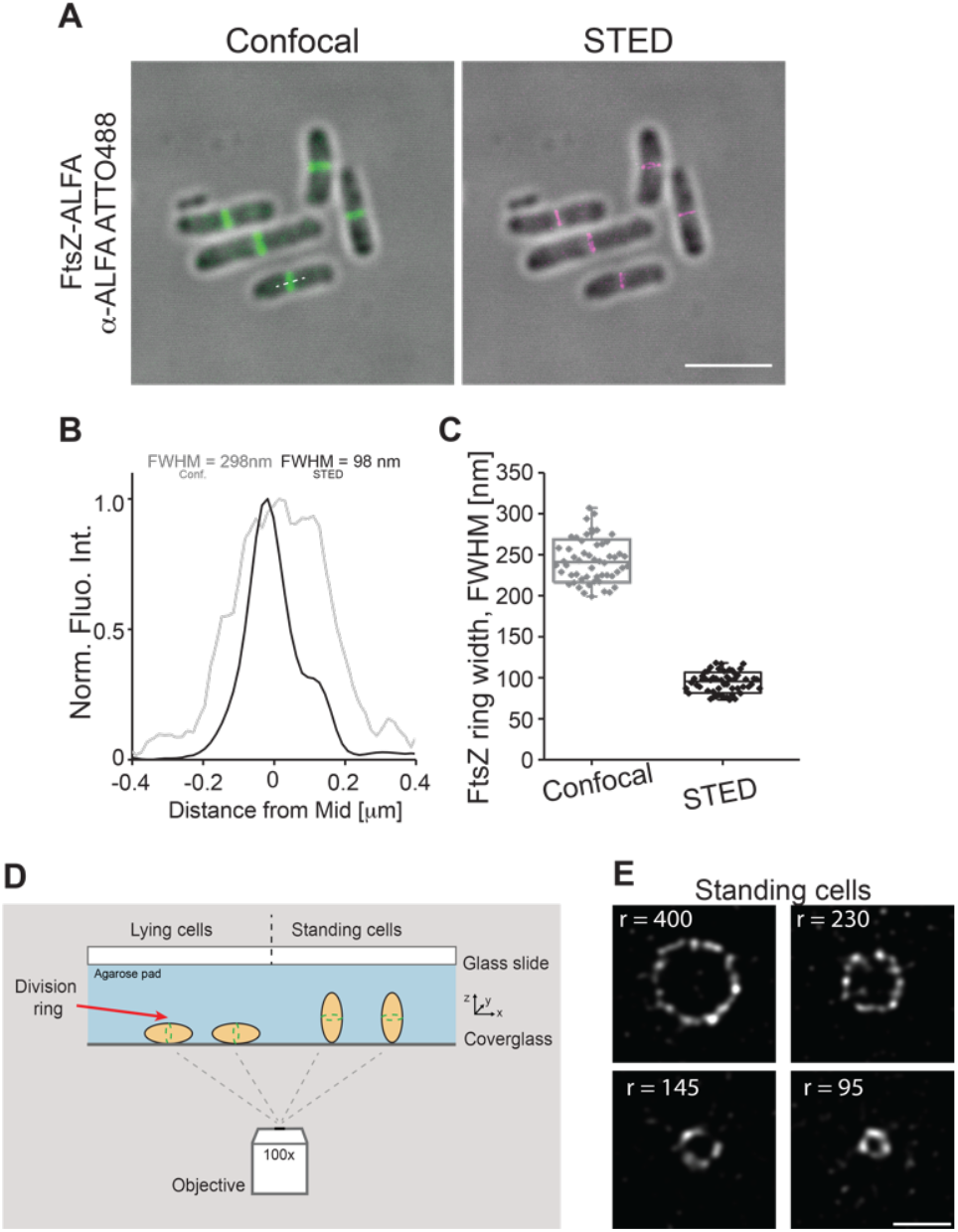
Super-resolution STED imaging of FtsZ-ALFA using an α-ALFA nanobody labelled with ATTO488. **A**, Fixed cells immunolabelled and imaged using confocal and STED microscopy. Fluorescence signal overlayed with bright field images. Scale bar = 4 μm. **B**, Longitudinal width (Full Width of Half Maxima) of a representative FtsZ ring. Conf = Confocal. **C**, Apparent widths (FWHM) were extracted from line scans as indicated by the white dotted line in the lower cells in the confocal image (A). The mean widths were 243 ± 26 nm (confocal) and 93 ± 12 nm (STED) (n = 54). Boxes indicate S.D., midline is the mean and whiskers incorporate 1-99% of the data range. **D**, Schematic representation of cells lying on an agarose pad and trapped in a vertical position. **E**, STED imaging of FtsZ-ALFA cells labelled with α-ALFA-ATTO488 trapped in a vertical position in micron holes. Representative FtsZ rings of various radii. r indicate radius. Scale bar = 500 nm.

### Live cell PALM using intracellular expressed fluorescent nanobodies

A limitation of the immunofluorescence approach is that the cells were fixed, so it was not possible to follow the dynamics of FtsZ-ALFA over time. To circumvent this problem, we transformed the FtsZ-ALFA strain with a plasmid encoding for the α-ALFA nanobody fused to the photoconvertible mEos3.2 (a FP compatible with single-molecule imaging). The main benefits with this approach are that (1) we could control the amount of mEos3.2-α-ALFA (so that not all FtsZ-ALFA molecules were labelled) and (2) we could produce mEos3.2-α-ALFA when required. In PALM the localization of single molecules from stochastically photoswitched or photoactivated fluorescent proteins are determined at sub-pixel resolution in a large series of sequential images from which a composite super-resolution image is created (Coltharp and Xiao 2012, Lelek, et al. 2021). After titrating the amount of mEos3.2-α-ALFA it was possible to identify conditions, where our experimental single-molecule approach showed clear FtsZ accumulation at midcell without affecting cell morphology or growth (Figure 3 and 4A).

**Figure 3.**
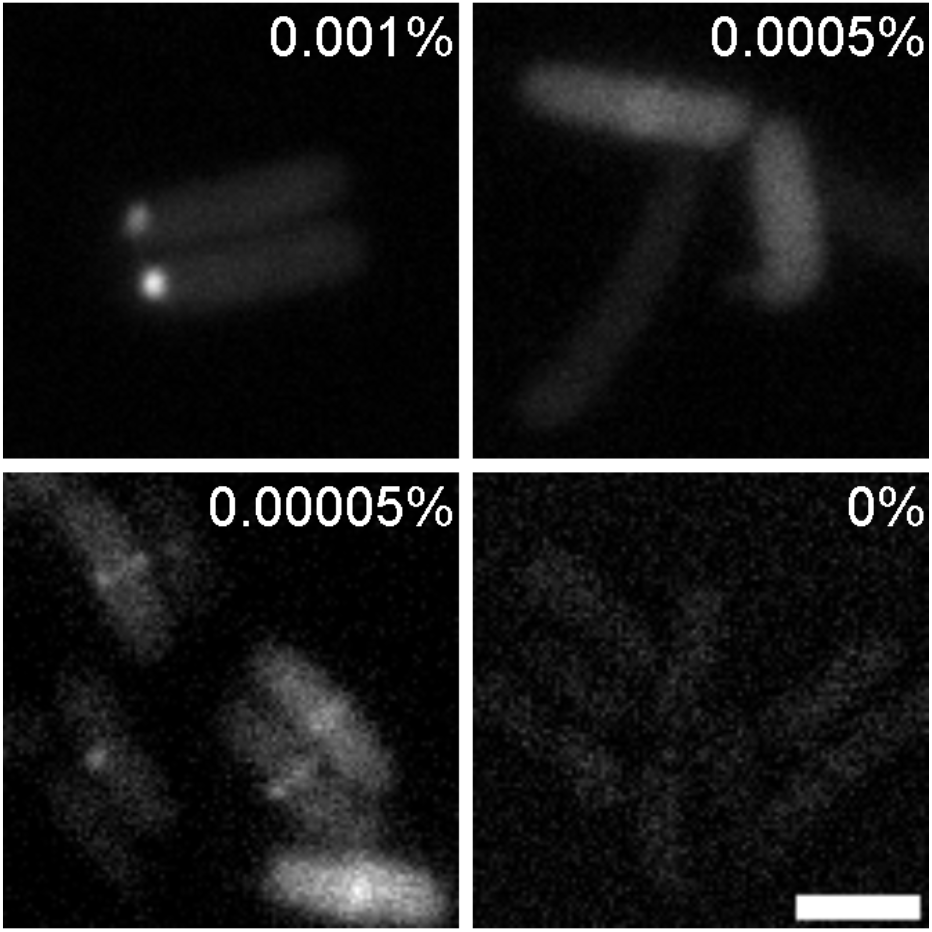
Efficient FtsZ-ALFA/mEos3.2-α-ALFA ring detection is concentration dependent. Epifluorescnece imaging on FtsZ-ALFA tagged with mEos3.2-α-ALFA in live E. coli cells. mEos3.2 was excited and captured in the green state using a 488-laser line. Images of various conditions with different Arabinose concentrations, ranging from 0.001% to 0%. Too high induction (0.001% and above) produced large bright protein aggregates at the poles, 0.0005% produced too high background, while no induction resulted in essentially no fluorescence signal. As clearly can be seen, under our experimental conditions, 0.00005% Arabinose gave best results. Scale bar = 2 μm.

**Figure 4.**
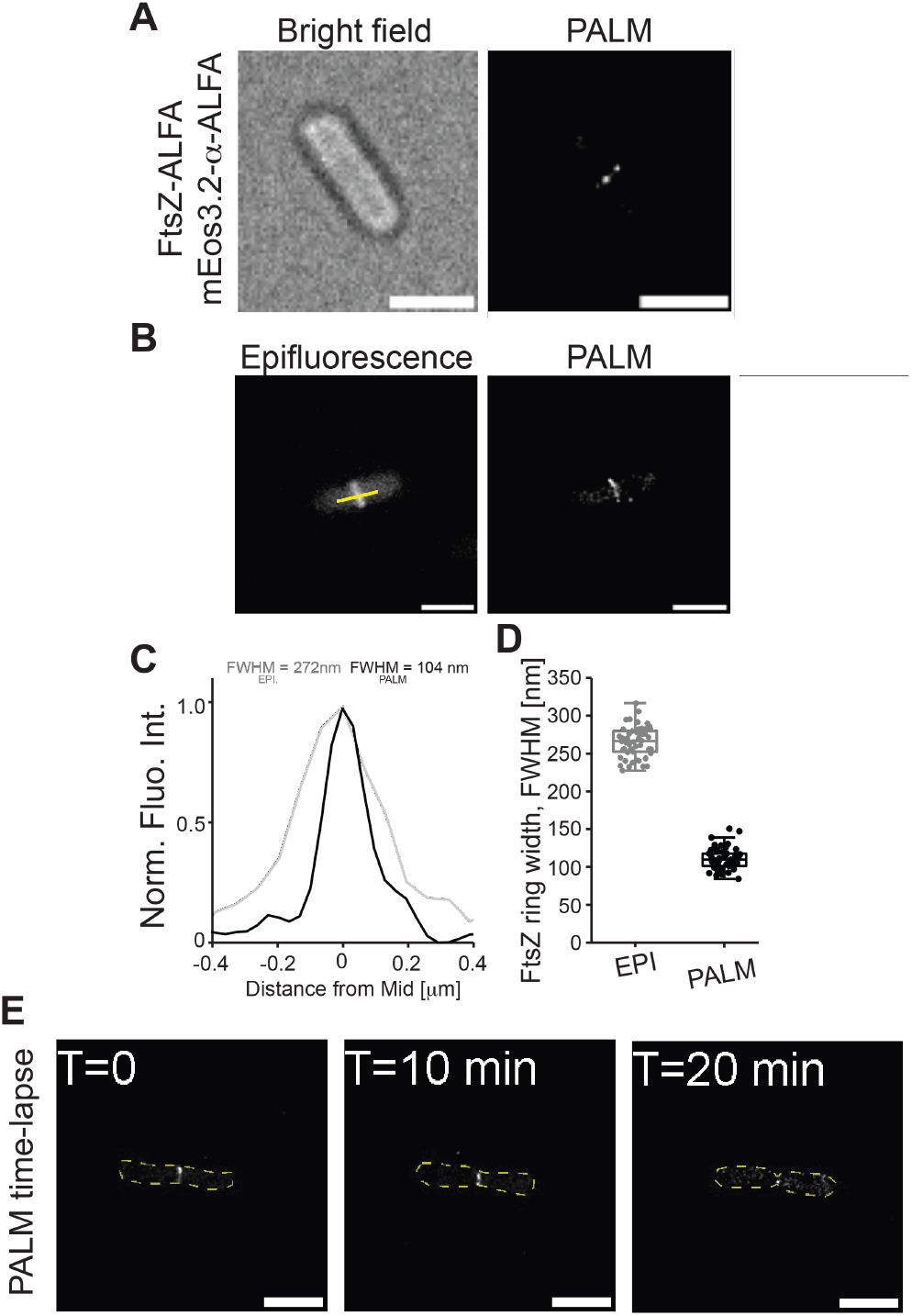
Live cell single-molecule PALM imaging of FtsZ-ALFA using a co-expressed mEos3.2-α-ALFA nanobody. **A**, A typical live cell imaged by bright field optics and using PALM. **B**, Comparison of a FtsZ-ALFA ring using Epifluorescence and PALM. **C**, Longitudinal width (Full Width of Half Maxima, FWHM) of a representative FtsZ-ALFA ring as determined by fitting a gaussian to fluorescence traces generated drawn over the FtsZ-ALFA rings (as in (B)). EPI = Epifluorescnece. **D**, Apparent widths (FWHM) from line scans as indicated by the yellow line in the lower cells in the epifluorescence image (B). The mean widths were 266 ± 22 nm (epifluorescence) and 110 ±11 nm (PALM) (n = 109). Boxes indicate S.D., midline is the mean and whiskers incorporate 1 - 95% of the data range. **E**, Images from a typical time-lapse PALM sequence. Z-ring radius is clearly decreasing over time. Scale bars = 2 μm.

Again, quantitative measurements of the Full Width at Half Maxima (FWHM) indicated a noticeable resolution improvement over diffraction limited imaging (Figure 4B-C). These Z-rings had an average lateral width of 110 ± 11 nm (n = 109) (Figure 4C), again similar to previously reported values using live cell PALM (Coltharp, et al. 2016, Fu, et al. 2010). One important consideration with this approach is that the expression of mEos3.2-α-ALFA needs to be optimised by titration. If expression is too low it will not be possible to detect the ALFA nanotag, and if the expression is too high the background fluorescence will be too high (as the mEos3.2-α-ALFA is fluorescent even when it is not bound to the ALFA nanotag) (Figure 3) (de Beer and Giepmans 2020). This latter problem is unique to the live cell imaging as it is not possible to wash away unbound mEos3.2-α-ALFA (as it is with IFM approaches).

The main advantage of this approach over IFM approaches is arguably that cells are imaged live and thus can dynamic processes be followed in super-resolution. To capture FtsZ dynamics at single-molecule resolution, we imaged cells on agarose pads over time. We acquired images every 10 minutes as it was not feasible to decrease the interwall between imaging due to the limited pool of mEos3.2-α-ALFA molecules. Time-lapse imaging showed dynamic FtsZ-rings with decreasing radius over time (Figure 4D).

## Conclusions

In summary, here we have demonstrated that nanotag/nanobody labelling approaches can be utilized to image bacterial cell division proteins at super resolution. Initially, FtsZ-ALFA rings were visualised in fixed cells using an immunofluorescence microscopy (IFM) protocol and super-resolution STED imaging (see also (Söderström, et al. 2018a, Söderström, et al. 2018b, Strauss, et al. 2012)).

In a second step, we also demonstrated the feasibility of nanobody labelling in live bacteria for the first time. We used live cells in a single-molecule PALM imaging approach by introducing plasmid encoded α-ALFA nanobodies fused to a fluorescent protein (mEos3.2). This approach allowed protein specific cell division dynamics at single-molecule resolution to be monitored over time. The advantages of our live cell nanobody approach over conventional fluorescent protein fusions are two-fold: the ALFA-tags are small and are therefore less likely to interfere with normal protein function, and the generation of the nanobody itself is controllable by induction meaning that its production can be restricted to biologically relevant time-points. We believe that this can easily be translated to a wide range of other events in bacteria and aid others looking to study time dependent processes at single-molecule resolution in systems otherwise challenging to label using standard approaches.

## Methods

### Strain engineering

The FtsZ-ALFA strain was made using the CRMAGE method (Ronda, et al. 2016). In short, the MG1655 strain was transformed with the pMA7CR.2.0 (amp^R^) and pZS4int-tetR (spec^R^) plasmids, then grown overnight at 37°C (with shaking) in LB Lennox media containing 100 μg/mL ampicillin and 50 μg/mL spectinomycin. The culture was back-diluted 1:100 and grown to an OD_600_ of 0.4-0.5. Expression of the λ RED β-protein was induced with 0.2% (w/v) L-arabinose for 15 min. Cells were washed three times in ice-cold water and resuspended in 400 μL of ice-cold water. A 50μL aliquot of the cells was mixed with 1 μL of 5 μM CRMAGE oligo (Integrated DNA Technologies, USA) and 250 ng of the pMAZ-SK (kana^R^) plasmid (containing the sgRNA for the native *ftsZ* locus) and transferred to an electroporation cuvette with a 1 mm gap. Cells were electroporated with 1.8kV and then mixed with 950 μL of LB media containing 100 μg/mL ampicillin and 50 μg/mL spectinomycin. After a 1-hour recovery at 37°C with shaking, 50 μg/mL of kanamycin was added, and cells were incubated for 2 hours at 37°C. The Cas9 protein was induced by the addition of 200 ng/mL aTetracycline and the culture was further incubated at 37°C (with shaking) overnight. A dilution series was plated on LB agar containing 100 μg/mL ampicillin, 50 μg/mL spectinomycin and 50 μg/mL of kanamycin. Colonies were screened by diagnostic PCR to identify the strain. The sequence of the CRMAGE oligo is listed in Supplementary Table S1. The pMAZ-SK plasmid containing the sgRNA was constructed using the *in vivo* cloning approach (Bubeck, et al. 1993, Garcia-Nafria, et al. 2016, Jones and Howard 1991). In short, the plasmid backbone was amplified by PCR using primers P9 and P10 and the Q5 DNA polymerase (New England Biolabs, USA, NEB #M0491). These primers were obtained from Eurofins Genomics (Germany) and are listed in Supplementary Table S1. The plasmid was sequenced verified by Eurofins Genomics (Germany).

The engineered FtsZ-ALFA strain was subsequently cured from the pMAZ-SK plasmid as previously described with minor modifications (Ronda, et al. 2016). In short: Cells were grown overnight then washed twice with fresh LB and back diluted in fresh LB with aTetracycline 200 ng/mL and 0.2% (w/v) of L-Rhamnose and grown for 24 h. Cultures were plated on Amp/Spect (100 μ g/mL ampicillin, 50 μ g/mL spectinomycin) and Kan plates (50 μ g/mL Kanamycin). The final cured strain MG1655 (FtsZ-ALFA(G55:Q56)) was denoted AB003.

### Plasmid construction

The expression plasmid pBAD/HisB-mEos3.2-ALFA (kanaR) was constructed using the pBAD/HisB(TIR^EVOL^)-mEos3.2 (ampR) backbone (Addgene plasmid #189720; (Shilling, et al. 2022) ref). Initially pBAD/HisB(TIR^EVOL^)-mEos3.2 (ampR) was linearised by PCR using the primers P1 and P2. The coding sequence for the α-ALFA NB was amplified by PCR from the mEGFP-NbALFA plasmid (Addgene plasmid #159986; (Jin, et al. 2021)) using the primers P3 and P4. This PCR product contained 5’ and 3’ extensions (19 and 18 bp’s, respectively) that overlapped with the desired integration site in the pBAD/HisB(TIR^EVOL^)-mEos3.2 (ampR) backbone so that they could be ligated using the *in vivo* cloning approach (Bubeck, et al. 1993, Garcia-Nafria, et al. 2016, Jones and Howard 1991). In short, this required that the PCR products were treated with DpnI for 60 minutes at 37 ^o^C, then transformed into MC1061 strain using a standard heat shock protocol. The final clone was called pBAD/HisB-mEos3.2-

ALFA (ampR). In a second step, the Tn3.17 fragment containing the coding sequence for ß-lactamase (and conferring resistance to ß-lactam antibiotics; ampR) was replaced with the Tn903.1 fragment containing the aminoglycoside 3’ phosphatase (and conferring resistance to aminoglycoside antibiotics; kanaR). pBAD/HisB-mEos3.2-ALFA (ampR) was linearised by PCR using the primers P5 and P6. The coding sequence for the Tn903.1 was amplified by PCR from the pET28 (T7p^CONS^/TIR2)-sfGFP plasmid (Addgene plasmid #154464;(Shilling, et al. 2020)) using the primers P7 and P8. This PCR product contained 5’ and 3’ extensions (15 and 18 bp’s, respectively) that overlapped with the integration site in the pBAD/HisB-mEos3.2-ALFA backbone so that they could be ligated using the *in vivo* cloning approach (Bubeck, et al. 1993, Garcia-Nafria, et al. 2016, Jones and Howard 1991). The final clone was called pBAD/HisB-mEos3.2-ALFA (kanaR). PCR was carried out with the Q5 DNA polymerase (New England Biolabs, USA, NEB #M0491). Primers were obtained from Eurofins Genomics (Germany) and are listed in Supplementary Table S1. Plasmids were sequenced verified by Eurofins Genomics (Germany).

### Bacterial growth and plasmid expression

Pre-cultures of AB003 with or without pBAD/HisB-mEos3.2-ALFA were grown overnight in 20 ml of LB with appropriate antibiotics (Ampicillin 100 μg ml^-1^, Kanamycin 50 μg ml^-1^) at 30 °C. The following morning the cultures were back-diluted 1:50 in LB incubated at 30 °C to OD_600_ 0.3 (~ 2 h). AB003 without the plasmid were processed for immunofluorescence and STED (see ‘*Cell fixation and nanobody labelling for STED’*). Cultures of AB003 transformed with pBAD/HisB-mEos3.2-ALFA were pelleted and resuspended in fresh M9 minimal media supplemented with 0.2% casaminoacids, 1 mM MgSO4, and 0.00005% arabinose (to initiate production mEos3.2-α-ALFA production) and grown at 30 °C for an additional two hours.

Four μl of cell culture was added to an agarose pad and taken for live cell imaging.

### Cell fixation and nanobody labelling for STED

Cells were fixed and immunodecorated as described previously (Söderström, et al. 2018b). Briefly, 900 μl of ice-cold methanol was added to 100 μl of cell culture (OD_600_ ~ 0.2 – 0.5) and incubated for 5 minutes on ice. Cells were harvested by centrifugation and resuspended in 100 μl of 90 % (v/v) ice-cold methanol. For nanobody labelling of cells, roughly 20 μl of cell suspension applied on a cover glass coated with Poly-L-Lysine or on agarose micron-hole beds and left for ~ 2 minutes before excess culture was removed. After drying, the cells were treated with lysozyme solution (0.2 mg/mL lysozyme, 25 mM Tris-HCl pH 8.0, 50 mM glucose, 10 mM EDTA) for 7-8 min and then rinsed with PBSTS [140 mM NaCl, 2 mM KCl, 8 mM Na_2_HPO_4_, 0.05% (v/v) Tween 20, 20% (w/v) sucrose], before they were treated with methanol for 1 min and acetone for 1 min and dried. Fixed cells were incubated with 3% (w/v) bovine serum albumin in PBSTS for at least 15 min. Thereafter the cells were washed with PBSTS and incubated with fluorescently labelled nanobodies *(i.e*. α-ALFA-ATTO488 and α-ALFA-ATTO647, gift from Dr. Hansjörg Götzke (NanobioTechnologies)) for 2-3 hours at RT. Cells on cover glasses were placed on glass slides with a drop of ProLong Gold antifade as mounting medium and left overnight to harden before imaging, while cells in agarose micron-hole beds were covered with a pre-cleaned cover glass.

### Imaging

STED images were acquired on a Leica TCS SP8 STED 3X system, using a HC PL Apo 100x oil immersion objective with NA 1.40. Fluorophores were excited using a white excitation laser operated at 488 nm (ATTO488), a STED laser line operated at 592 nm, and detection time-delay of 0.9 - 2.5 ns. The total depletion laser intensity was on the order of 20 - 500 MW/cm^2^ for all experiments. The final pixel size was 13 nm and scanning speed 600 Hz. The pinhole size was varied between 0.4-0.9 AU; a smaller pin-hole when imaging cells standing trapped in micron holes and larger when imaging cells lying flat on agarose pads.

Live cell PALM imaging was performed on a Nikon (Ti2-E) N-STORM v5 with NIS v.5.30 using a 100x 1.49 NA oil objective in semi-TIRF mode. A stage-top environmental chamber (Oko lab) was used for temperature control, imaging was performed at 37 °C for live cell and time-lapse imaging. Cover glass slides were washed with 80% EtOH, air dried, cleaned for at least 3 minutes with a plasma cleaner (Harrick plasma, PDC-23G) and used within 15 minutes of cleaning. Prior to single-molecule acquisition, the green state of mEos3.2 was excited by a 488 nm laser to acquire an epifluorescence image using a FITC emission filter cube. For single-molecule acquisition, mEos3.2 was photoconverted to its red state continuously by a 405 nm laser with increasing working powers ranging between 0.1 and 5 W/cm^2^. As readout, mEos3.2 was excited by a 561 nm laser line operating at an average power between 1 and 2 kW/cm^2^. The emission was collected by a quad band (Quad405/488/561/647 filter dual cSTORM). The exposure time was 20 ms and ~ 3000 - 4000 images were typically acquired for each set of images. Drift correction during image acquisition was minimised using the integrated PFS4 (Perfect Focus System). Images were captured using a sCMOS Flash 4.0 v3 (Hamamatsu) camera.

### Image analysis

STED images were deconvolved using Huygens Professional deconvolution software (SVI, the Netherlands). The resolution in our STED images was ~ 50 nm (Söderström, et al. 2018b). Single-molecule raw data were analysed using the Fiji plug-in ThunderSTORM (Ovesny, et al. 2014). Super resolution images were rendered using Gaussian blur of 20 nm for visualization. The relatively short PALM image acquisition time (50 Hz) combined with that the overall system drift was consistently less than one half pixel or less over the series acquisition time, therefore were no fiducial markers needed (Supplementary Figure 1). Z-ring widths in cells lying down were extracted from line traces of the midcell. A Gaussian distribution was fitted to the intensity profiles in order to extract the Full Width at Half Maximum (FWHM). Radii from cells trapped standing were estimated by manually fitting a circle over the fluorescence traces in Fiji. Statistics and graphs were generated using custom Matlab scripts, Origin9 Pro and PlotsOfData (Postma and Goedhart 2019)

### Cell length measurements

Cells of WT, FtsZ-ALFA and FtsZ-ALFA/mEos3.2-α-ALFA were harvested from growth cultures by centrifugation. An aliquot (2 μl) was placed on an agarose (1.5% w/w) bed and directly imaged under bright-field illumination. Cell lengths of at least 200 cells from each strain were determined using MicrobeJ (Ducret, et al. 2016). Statistics and graphs were generated using custom Matlab scripts, Origin9 Pro and PlotsOfData (Postma and Goedhart 2019).

### Western blotting

Cell extracts from a volume corresponding to 0.2 DO_600_ units were collected for different strains. The extracts were suspended in loading buffer and resolved by sodium dodecyl sulphate polyacrylamide gel electrophoresis (SDS-PAGE). Proteins were transferred to nitrocellulose membranes using a semi-dry Turbo Transfer-Blot apparatus (Bio-Rad). The membranes were blocked in 5 % (w/v) milk and probed with antisera to FtsZ [1:4000] (Agrisera, Sweden).

## Data availability

The data supporting the findings of the study are available in this article, or from the corresponding authors upon request.

## Acknowledgements

The authors would like to thank Alex Toftgaard Nielsen and Morten Nørholm (DTU, Denmark) for plasmids, and help setting up the CRMAGE method. We would also like to thank Dr. Hansjörg Götzke (NanobioTechnologies) for the fluorescently labelled nanobodies. This study was supported by a JSPS KAKENHI young scientist grant (JP17K15694, Japan), a CPDRF grant (UTS, Australia) and an ARC Discovery Project grant (DP220101143) to B.S. Work in the SCB unit at OIST is funded by core subsidy from Okinawa Institute of Science and Technology Graduate University. Hannah Brown (UTS) is acknowledged for help with the Western blotting. D.O.D. was supported by grants from Novo Nordisk Fonden (0071844) and Carl Trygger stiftelse (CTS 21:1637).

## Author contributions

B.S. and D.D. conceptualized the study and designed the experiments. B.S., A.B., E.W. and A.P. performed the experiments. H.C. contributed new reagents. All authors helped with data analysis. B.S and D.D. wrote the manuscript with input from all authors. All authors approved the final version of the manuscript.

## Competing interests

The authors declare no competing financial interests.

## Notes

### Competing Interest Statement

The authors have declared no competing interest.

