## Supplementary Info for "Application of nanotags and nanobodies for live cell single-molecule imaging of the Z-ring in *Escherichia coli*"

Emma Westlund^1, #^, Axel Bergenstråle ^2,#^, Alaska Pokhrel^1^,
Helena Chan^3^, Ulf Skoglund^3^, Daniel O. Daley^2,^* and Bill Söderström^1,^*

^1^ Australian Institute for Microbiology and Infection, University of Technology Sydney, Ultimo, NSW, 2007 Australia.

^2^ Department of Biochemistry and Biophysics, Stockholm University, SE-106 91 Stockholm, Sweden.

^3^ Structural Cellular Biology Unit, Okinawa Institute of Science and Technology, 904-0495 Okinawa, Japan^.^

\# equal contribution

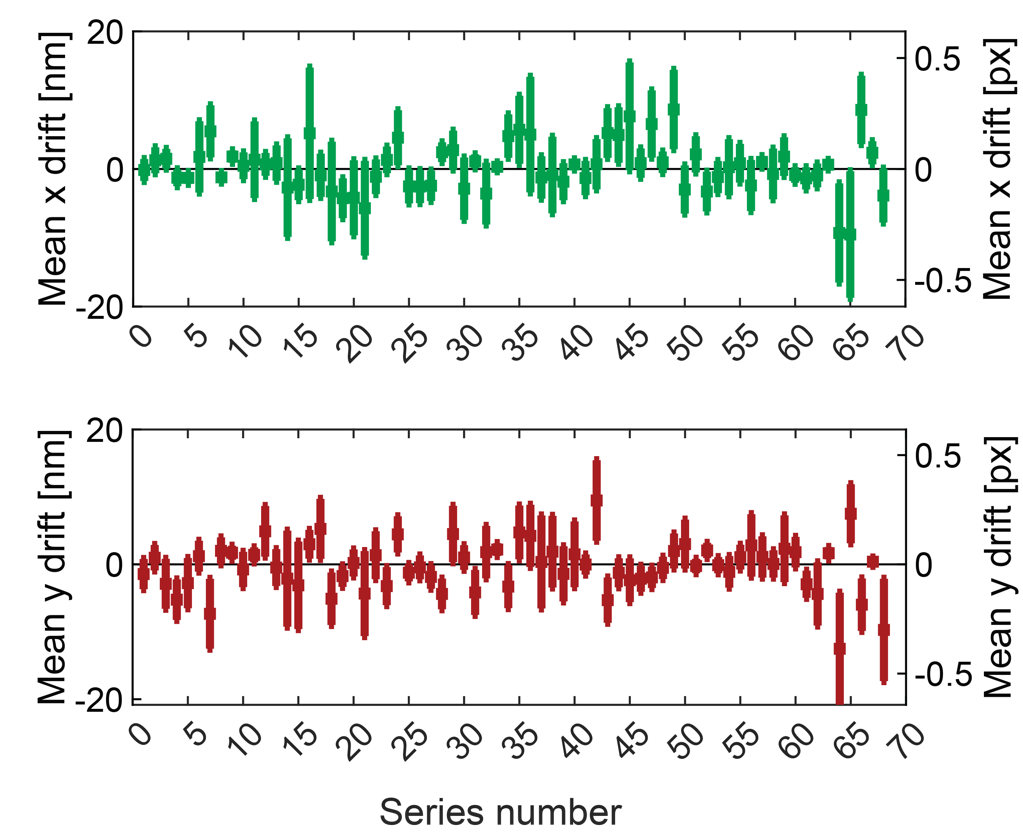

**Supplementary figure 1. Mean drift during single-molecule image acquisition.**

The mean drift ( ± standard deviation) over the full series in the x- and y-direction for randomly chosen image series during PALM imaging of mEos3.2. The drift was consistently less than one half pixel.

**Supplementary Table S1. Primers and CRMAGE oligonucleotide used.**

Primers

P1	pBAD.fwd	5’-GAATTCGAAGCTTGGCTGTTTTGGCGG-3’

P2	mEos.rev	5’-TCGTCTGGCATTGTCAGGCAATCC-3’

P3	NbALFA.fwd	5’-GCCTGACAATGCCAGACGAGGCAGTGGCAGTGGCAGTGGAG-3’

P4. NbALFA.rev	5’-CAGCCAAGCTTCGAATTCTTAGCTGCTCACAGTCACTTGGGTGCC-3’

P5	Bla loop out primer 1	5’-AGAGTTTGTAGAAACGCAAAAAG-3’

P6	Bla loop out primer 2	5’-CTGTCAGACCAAGTTTACTC-3’

P7	Aph intro primer 1	5’-GTTTCTACAAACTCTCATGAACAATAAAACTGTCTG -3

P8	Aph intro primer 2	5’-CTGACAGCTTAGAAAAACTCATCGAGCATCAAATG-3’

P9	FtsZ-crRNA-Fwd	5’- AGCGCTGCGTAAAACAGGTTTTAGAGCTAGAAATAGCAAG -3’

P10	FtsZ-crRNA-Rev	5’- TTTACGCAGCGCTTGTGTGCTCAGTATCTCTATCACTG -3’

CRMAGE oligo

NT-L-FtsZ	To insert ALFA nanotag at G55:Q56 with flanking sequences (GSTLE and LEGST)

5’GCGCCAGCGCCCAGTCCTTTGGTGATACCGCTACCGATTTGAATCGTCTGGGTGCTACCTTCCAGTGCCGTCAGACGACGACGCAGTTCCTGTTCCAGGCCGGATTCCAGGGTGCTACCTCCAACTGCTGTTTTACGCAGCGCTTGTGCATCGGTATTTACCGCGAAGA -3’
